# Social odors drive hippocampal CA2 place cell responses to social stimuli

**DOI:** 10.1101/2024.07.16.603738

**Authors:** Emma Robson, Margaret M. Donahue, Alexandra J. Mably, Peyton G. Demetrovich, Lauren T. Hewitt, Laura Lee Colgin

**Affiliations:** Center for Learning and Memory, The University of Texas at Austin, Austin, TX 78712; Department of Neuroscience, The University of Texas at Austin, Austin, TX 78712; Institute for Neuroscience, The University of Texas at Austin, Austin, TX 78712

**Keywords:** hippocampus, place cells, CA2, social cognition, social memory, social odor

## Abstract

Hippocampal region CA2 is essential for social memory processing. Interaction with social stimuli induces changes in CA2 place cell firing during active exploration and sharp wave-ripples during rest following a social interaction. However, it is unknown whether these changes in firing patterns are caused by integration of multimodal social stimuli or by a specific sensory modality associated with a social interaction. Rodents rely heavily on chemosensory cues in the form of olfactory signals for social recognition processes. To determine the extent to which social olfactory signals contribute to CA2 place cell responses to social stimuli, we recorded CA2 place cells in rats freely exploring environments containing stimuli that included or lacked olfactory content. We found that CA2 place cell firing patterns significantly changed only when social odors were prominent. Also, place cells that increased their firing in the presence of social odors alone preferentially increased their firing during subsequent sharp wave-ripples. Our results suggest that social olfactory cues are essential for changing CA2 place cell firing patterns during and after social interactions. These results support prior work suggesting CA2 performs social functions and shed light on processes underlying CA2 responses to social stimuli.

## 1. Introduction

Social recognition memory, or the ability to recognize and remember conspecifics, facilitates social interactions that are essential for animals to survive. Rats can discriminate between familiar conspecifics (Husted and McKenna, 1966; Thor and Holloway, 1982) based on their distinct characteristics and olfactory signatures (Gheusi et al., 1997; Popik et al., 1991; Sawyer et al., 1984). Many recent studies have implicated hippocampal area CA2 in social memory (for a review, see Oliva, 2022). Lesioning or inactivating CA2 pyramidal neurons leads to impaired social recognition memory in mice (Hitti and Siegelbaum, 2014; Stevenson and Caldwell, 2014) and optogenetic silencing of CA2 activity shows that CA2 is crucial for encoding, consolidating, and recalling social memories (Meira et al., 2018). Moreover, CA2 neurons are uniquely enriched in receptors selective for a variety of social neuropeptides, including vasopressin and oxytocin (Lee et al., 2008; Lin et al., 2018; Pagani et al., 2015; Raam et al., 2017; Wersinger et al., 2002, 2008).

All subregions of the hippocampus contain place cells, neurons that are selectively activated in a particular location in space (their “place fields”) (O’Keefe, 1976; O’Keefe and Dostrovsky, 1971). Place cells alter their firing in response to significant environmental changes in a process called “remapping” (for a review, see Colgin et al., 2008). During “global” remapping, a place cell’s place field may appear, disappear, or change location. During “rate” remapping, a place cell’s place field location does not move, but its in-field firing rate changes. Neurophysiological recordings of place cells have shown that place fields in hippocampal subregions CA1 and CA3 remain relatively stable in unchanging environments (Muller et al., 1987; Thompson and Best, 1990). In contrast, CA2 place cells show gradual changes in firing patterns over time (Alexander et al., 2016; Mankin et al., 2015). CA2 place cells also appear to be highly sensitive to small updates to familiar environments (Wintzer et al., 2014). Previous work has shown that a significant proportion of CA2 place cells globally remap when a familiar conspecific is presented (Alexander et al., 2016). This remapping in response to social stimuli may support the ability of CA2 to encode social memories (Hitti and Siegelbaum, 2014). However, it remains unclear whether CA2 place field changes that occur in response to the presentation of a conspecific rat are caused by integration of multimodal social stimuli or primarily due to a specific sensory modality associated with the social interaction.

Previous work has also shown that CA2 firing patterns during sleep are altered following a social experience (Oliva et al., 2020). During awake rest or sleep, hippocampal place cells fire during distinctive events in the hippocampal local field potential (LFP) known as sharp wave-ripples. Place cells that were active during exploratory behaviors later reactivate during sharp wave-ripples, and reactivated place cell ensembles are believed to support memory consolidation (Ego-Stengel and Wilson, 2010; Ramadan et al., 2009; Wilson and McNaughton, 1994). CA2 place cells that represent a conspecific have been shown to reactivate during sharp wave-ripples following a social experience (Oliva et al., 2020). Further, disrupting CA2 sharp wave-ripples impairs social recognition memory (Oliva et al., 2020). Therefore, the reactivation of CA2 place cells that encode a social experience may promote consolidation of social memories (Oliva et al., 2023).

In this study, we recorded CA2 place cells in dorsal hippocampus of rats during free exploration of a two-dimensional spatial environment and during subsequent rest periods. We compared changes in CA2 place cell firing patterns across sessions in which different social or non-social stimuli spanning various sensory modalities were presented. Significant global remapping was observed in CA2 place cells when social odors were presented in the absence of a rat and when a familiar rat was presented in a soiled home cage containing social odors (as in our prior study; Alexander et al., 2016). However, no significant changes in CA2 place cell firing rate maps were observed when a familiar rat was presented in a clean and relatively odorless cage or when a non-social odor was presented. These results suggest that olfactory cues are the key sensory component of social experiences that drives CA2 place cell remapping. Furthermore, CA2 place cells that increased their firing rates in response to social odors preferentially increased their firing rates in sharp wave-ripples during subsequent rest periods. These results improve our understanding of how CA2 place cells code information related to social interactions.

## 2. Materials and Methods

### 2.1. Subjects

Six wild-type male Sprague-Dawley rats (Inotiv, USA) were used for this study. Rats were between the ages of 4 and 10 months at the time of surgery. Before surgery, rats were double or triple housed and were pre-trained to freely explore an open field enclosure. After surgery, rats were singly housed in custom-built acrylic cages (40 cm x 40 cm x 40 cm) containing enrichment material (wooden blocks, paper towel rolls, etc.) and maintained on a reverse light cycle (light: 8 p.m. – 8 a.m.). Rats were housed next to their former cage mates after recovering from surgery and throughout behavioral testing. Rats recovered from surgery for at least one week before behavioral training resumed. All behavioral experiments were performed during the dark cycle. In order to encourage spatial exploration, four of the six rats were placed on a food deprivation regimen. All rats maintained ∼98% of their free-feeding body weight throughout the experiment. All experiments were conducted according to the guidelines of the United States National Institutes of Health Guide for the Care and Use of Laboratory Animals and under a protocol approved by the University of Texas at Austin Institutional Animal Care and Use Committee.

### 2.2. Surgery and tetrode positioning

Drives with 14 independently movable tetrodes were implanted in five of the rats. A drive with 21 independently movable tetrodes was implanted in one of the rats. Drives were implanted above a relatively lateral part of right dorsal hippocampus (anterior-posterior −3.8 mm from bregma, medial-lateral −3.0 mm from bregma). Because the diameter of the bundle of cannula surrounding tetrodes was wider than the mediolateral extent of CA2, this placement ensured that at least some of the tetrodes would hit CA2. To stabilize the recording drives, eleven bone screws were affixed to the skull and covered in dental acrylic. Two of the screws were connected to the recording drive ground. Prior to surgery, tetrodes were built from 17 μm polyimide-coated platinum-iridium (90/10%) wire (California Fine Wire, Grover Beach, California, USA). The tips of tetrodes designated for single-unit recording were plated with platinum to reduce single-channel impedances to ∼150 to 300 kOhms. All tetrodes were lowered ∼1 mm on the day of surgery. Thereafter, tetrodes were slowly lowered to the hippocampal pyramidal cell body layer over the course of several weeks.

One tetrode was designated for use as a reference for differential recording. All four wires of this tetrode were connected to a single channel on the electrode interface board. The differential recording reference tetrode was placed in an electrically quiet area approximately 1 mm above the hippocampus and adjusted as needed to ensure quiescence. The reference signal was duplicated using an MDR-50 breakout board (Neuralynx, Bozeman, MT, USA) and recorded continuously to ensure that unit activity or volume conducted signals of interest were not detected. Another tetrode was placed in the apical dendritic layer of CA1 to monitor LFPs and guide placement of the other tetrodes using estimated depth and electrophysiological hallmarks of the hippocampus (for example, sharp wave-ripples).

### 2.3. Data acquisition

Data were acquired using a Digital Lynx system and Cheetah recording software (Neuralynx, Bozeman, MT, USA). The recording setup has been described in detail previously (Hsiao et al., 2016; Zheng et al., 2016). Briefly, LFP signals from one randomly chosen channel within each tetrode were continuously recorded at a 2000 Hz sampling rate and filtered in the 0.1–500 Hz band. Input amplitude ranges were adjusted before each recording session to maximize resolution without signal saturation. Input ranges for LFPs generally fell within ±2,000 to ±3,000 μV. To detect unit activity, all four channels within each tetrode were bandpass filtered from 600 to 6,000 Hz. Spikes were detected when the filtered continuous signal on one or more of the channels exceeded a threshold set daily by the experimenter, which ranged from 55–65 µV. Detected events were acquired with a 32,000 Hz sampling rate for 1 ms. For both LFPs and unit activity, signals were recorded differentially against a dedicated reference channel (see “Surgery and tetrode positioning” section above).

Videos of rats’ behavior were recorded through the Neuralynx system with a resolution of 720 × 480 pixels and a frame rate of 29.97 frames/s. Rat position and head direction were tracked via an array of red and green or red and blue light-emitting diodes (LEDs) on a HS-54 or HS-27 headstage (Neuralynx, Bozeman, MT, USA), respectively.

### 2.4. Behavioral task

Rats were familiarized to an open field arena (1 m x 1 m with 0.5 m wall height) for a minimum of three days before recording started. Rats freely explored the open field arena for four 20-minute sessions per day, with 10-minute rest sessions preceding and following each active exploration session. During active exploration, small pieces of sweetened cereal or cookies were randomly scattered to encourage rats to explore the entirety of the arena. Rats had to cover at least 60% of the arena across each of the four sessions for a day to be included for further analysis. During each rest session, rats rested in a towel-lined, elevated flowerpot outside of the arena. A plexiglass standard rat housing cage was placed in one corner of the arena for all recording sessions. In the first and fourth sessions (A and A’), this cage contained only clean bedding. In the middle two sessions (B and B’) of the different experimental conditions, the cage contained various types of social stimuli (Figure 1). In the Social Odor condition, the soiled bedding from the former home cage of familiar rats (or a familiar rat) was used. In the Visual + Odor condition, a familiar rat was presented in its home cage containing soiled bedding. In the Visual condition, a familiar rat was placed in the stimulus cage with clean bedding and a filter-top lid to minimize social odors. In the Mirror condition, a familiar rat was placed in the stimulus cage with clean bedding and a filter-top lid, but the cage was lined with a one-way mirror attachment. This one-way mirror attachment prevented the stimulus rat from seeing the implanted rat, limiting visually driven reciprocal interactions between the two rats. In the Non-social Odor condition, three drops of hexyl acetate (Sigma-Aldrich Cat# 108154), which has a fruity smell, were applied to a wooden block which was placed underneath clean bedding in the stimulus cage. In the Control condition, a cage containing only clean bedding was presented in all four sessions.

**Figure 1.**
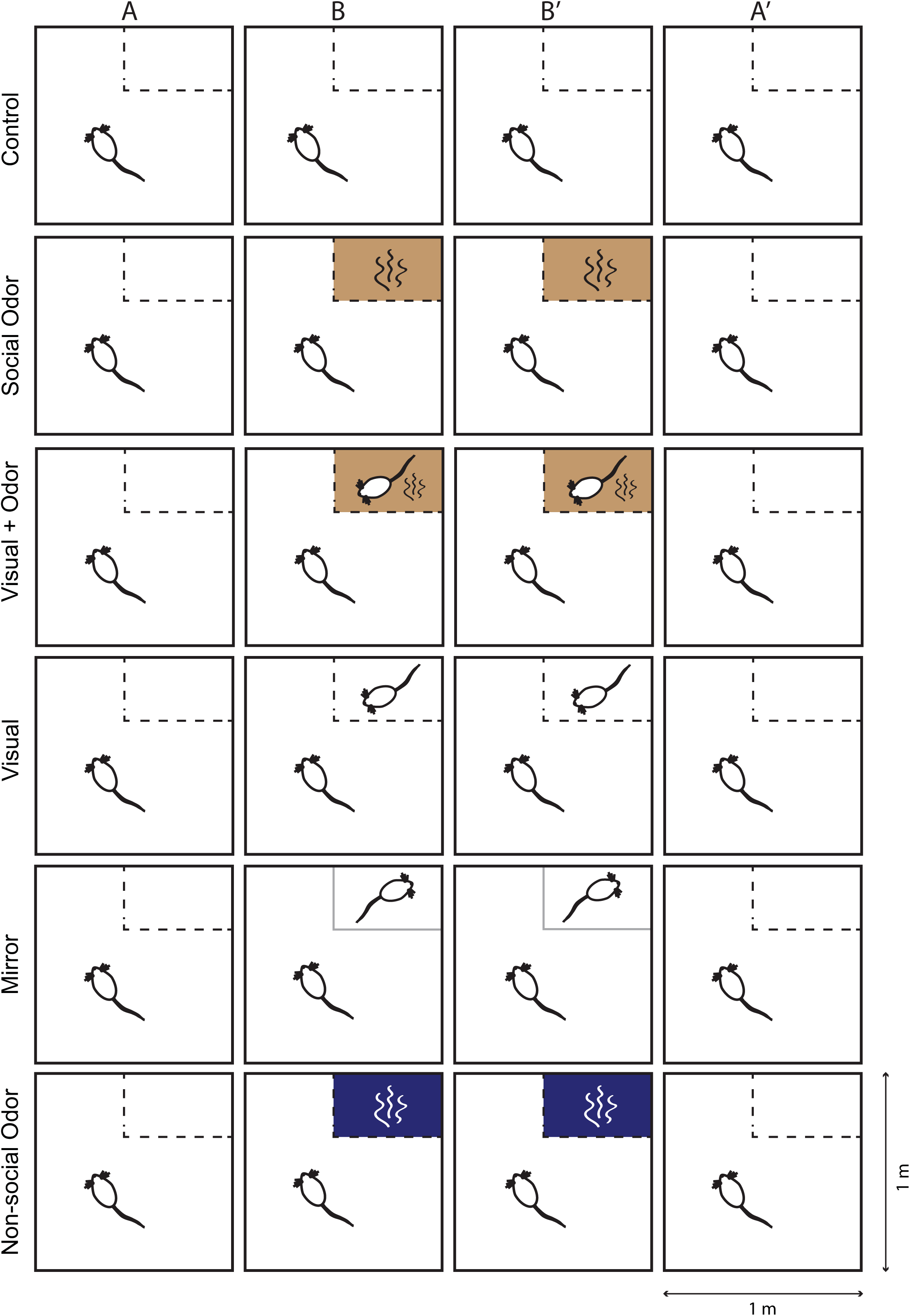
Behavioral task. Rats freely explored a familiar open field arena for four 20-minute sessions per day. A stimulus cage, a standard rat housing cage, was placed in a corner of the arena. In the first (A) and final (A’) sessions of experimental conditions (i.e., Social Odor, Visual + Odor, Visual, Mirror, and Non-social Odor), and in all four sessions of the Control condition, the cage contained only clean bedding. In the middle two sessions (B and B’) of experimental conditions, a cage containing stimuli (“stimulus cage”) was presented. In the Social Odor condition, a stimulus cage containing soiled bedding of a familiar rat or rats was presented. In the Visual + Odor condition, a stimulus cage containing a familiar rat and the rat’s soiled bedding was presented. In the Visual condition, a familiar rat was placed in the stimulus cage with clean bedding and a filter-top lid to minimize social odors. In the Mirror condition, a familiar rat was placed in the stimulus cage with clean bedding, and the cage was lined with a one-way mirror attachment. This prevented the stimulus rat from seeing the implanted rat, thereby limiting reciprocal social interactions. In the Non-social Odor condition, a wooden block infused with hexyl acetate, a fruity-smelling odorant, was placed in the stimulus cage under clean bedding.

### 2.5. Histology and tetrode localization

Following recording, rats were perfused with 4% paraformaldehyde solution in phosphate-buffered saline. Brains were cut coronally in 30 μm sections using a cryostat. For two rats, brain sections that contained tetrode tracks were immunostained with a monoclonal antibody specific for the CA2-expressed protein, STEP (Cell Signaling, 4817, 1:1000 dilution, raised in mouse). Sections were then incubated in an anti-mouse secondary antibody and developed with 3,3-diaminobenzidine (brown) (as in Alexander et al., 2016). Sections were then counter-stained with cresyl-violet (purple/blue) to indicate cell bodies (Supplementary Figure 1). Brain sections that did not contain tetrode tracks were stained with cresyl violet only. In order to locate CA2 in the sections with tetrode tracks, we examined STEP staining in the cell body layer and identified the portion of the hippocampus where the cell body layer increases in size to denote the boundaries of CA2 (as in Lu et al., 2015). For four rats, brains were immunostained for the CA2 marker Purkinje Cell Protein 4 (PCP4) (Figure 2). For this immunostaining protocol, sections were washed three times for 15 minutes in tris-buffered saline (TBS) followed by a 10-minute water wash. Sections were then permeabilized and washed for 15 minutes in TBS containing 0.3% Triton-X followed by three 10-minute washes in TBS. Sections were blocked for 30 minutes in 10% normal goat serum in TBS. Sections were incubated overnight with rabbit anti-PCP4 (1:200, Sigma-Aldrich Cat# HPA005792) diluted in TBS containing 0.05% Tween. The next day, sections were washed twice for 10 minutes in TBS and incubated overnight with secondary fluorescent antibody (in one rat: Alexa Flour™-488 anti-rabbit, Thermo Fisher Scientific; in three rats: Alexa Flour™-555 anti-rabbit, Thermo Fisher Scientific). All washes and incubations were performed at room temperature. Slides were mounted using DAPI Fluoromount-G (Fisher Scientific). Tetrode recording sites were identified by comparing locations across adjacent sections.

**Figure 2.**
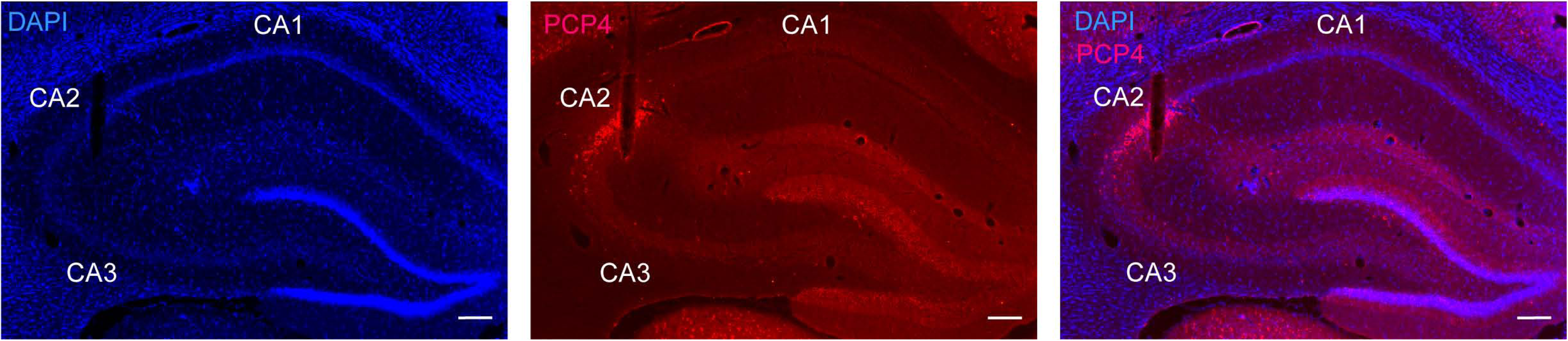
CA2 Histology. An example hippocampal section showing immunohistological identification of a tetrode track in CA2. To identify CA2, hippocampal sections were immunostained with a CA2 marker, Purkinje cell protein 4 (PCP4, red). DAPI nuclear staining is shown in blue. Scale bar, 200 µm.

### 2.6. Spike sorting and unit selection

Spike sorting was performed manually using graphical cluster-cutting software (MClust, A.D. Redish, University of Minnesota, Minneapolis, Minnesota) run in MATLAB (Mathworks). Spikes were sorted using two-dimensional representations of waveform properties (i.e., energy, peak, and peak-to-valley difference) from four channels. A single unit was accepted for further analysis if the associated cluster was well isolated from, and did not share spikes with, other clusters on the same tetrode. Units were also required to have a minimum 1 ms refractory period. Units with mean firing rates above 5 Hz were considered putative interneurons and excluded from further analysis. In order to be considered active in the arena, a unit had to reach a peak firing rate of at least 1 Hz. In order to be included in the sharp wave-ripple firing analysis, a unit had to have valid clusters in both the active exploration and rest sessions. CA2 cell yields for each condition are reported in Tables 1 and 2.

**Table 1.** Number of CA2 place cells recorded in each rat and each condition during active exploration sessions.

| Animal ID | Control | Social Odor | Visual +<br>Odor | Visual | Mirror | Non-social<br>Odor |
| --- | --- | --- | --- | --- | --- | --- |
| Rat 117 | 7 | 5 | 0 | 7 | 6 | 0 |
| Rat 122 | 51 | 32 | 0 | 38 | 49 | 0 |
| Rat 165 | 34 | 56 | 26 | 26 | 35 | 0 |
| Rat 256 | 28 | 13 | 13 | 13 | 21 | 11 |
| Rat 391 | 15 | 4 | 5 | 6 | 5 | 6 |
| Rat 418 | 20 | 35 | 41 | 18 | 0 | 29 |
| **Total** | **155** | **145** | **85** | **109** | **116** | **46** |

**Table 2.**
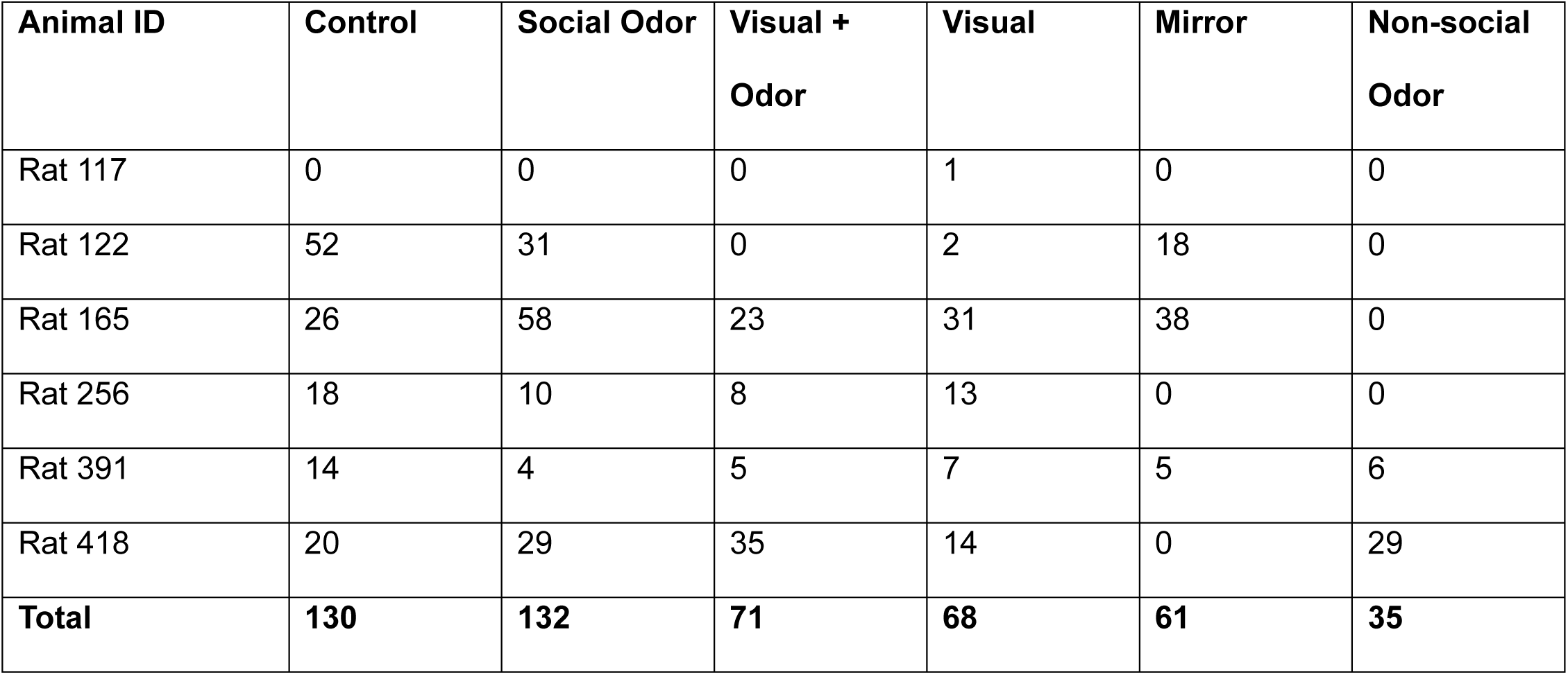
Number of CA2 place cells recorded in each rat and each condition during rest periods.

| Animal ID | Control | Social Odor | Visual +<br>Odor | Visual | Mirror | Non-social<br>Odor |
| --- | --- | --- | --- | --- | --- | --- |
| Rat 117 | 0 | 0 | 0 | 1 | 0 | 0 |
| Rat 122 | 52 | 31 | 0 | 2 | 18 | 0 |
| Rat 165 | 26 | 58 | 23 | 31 | 38 | 0 |
| Rat 256 | 18 | 10 | 8 | 13 | 0 | 0 |
| Rat 391 | 14 | 4 | 5 | 7 | 5 | 6 |
| Rat 418 | 20 | 29 | 35 | 14 | 0 | 29 |
| **Total** | **130** | **132** | **71** | **68** | **61** | **35** |

### 2.7. Place cell remapping analyses

Methods used to create firing rate maps for each single unit were based on methods used in our prior study (Alexander et al., 2016; example rate maps for the present study shown in Figure 3). First, the arena was divided into 4 cm^2^ bins. The number of spikes that occurred within each bin was divided by the time spent in that bin to determine the firing rate. Only spikes that occurred while the rat was traveling 5 cm/s or faster were included. The rate map was smoothed with a two-dimensional Gaussian kernel (standard deviation = 6 cm). To determine if a place cell remapped during presentation of a social stimulus, a Pearson correlation coefficient *R* was calculated for each unit between pairs of rate maps from control and social stimuli sessions (i.e., A-B, B-B’, B’-A’, A-B’, and A-A’ session pairs, see Figure 1). To determine if spatial correlation coefficients differed across session pairs and conditions (i.e., Social Odor, Visual + Odor, Visual, Mirror, Non-social Odor, Control), we used a generalized linear mixed model statistical analysis (IBM SPSS Statistics, version 29.0.2.0). Condition and session pair were fixed factors, session pairs were repeated measures within rats, and multiple place cells were nested within rats. A condition by session pair interaction effect was also included in the model to determine whether differences between session pairs varied across conditions. When a significant effect was observed, post-hoc pairwise comparisons were performed with Bonferroni correction for multiple comparisons. The estimated mean spatial correlations for each condition and session pair from the generalized linear mixed model are shown in Figure 4A. The estimated mean spatial correlations for each condition, collapsed across session pairs, are shown in Figure 4B. Individual spatial correlation values for each cell are shown for each session pair and each condition in Figure 4C-G.

**Figure 3.**
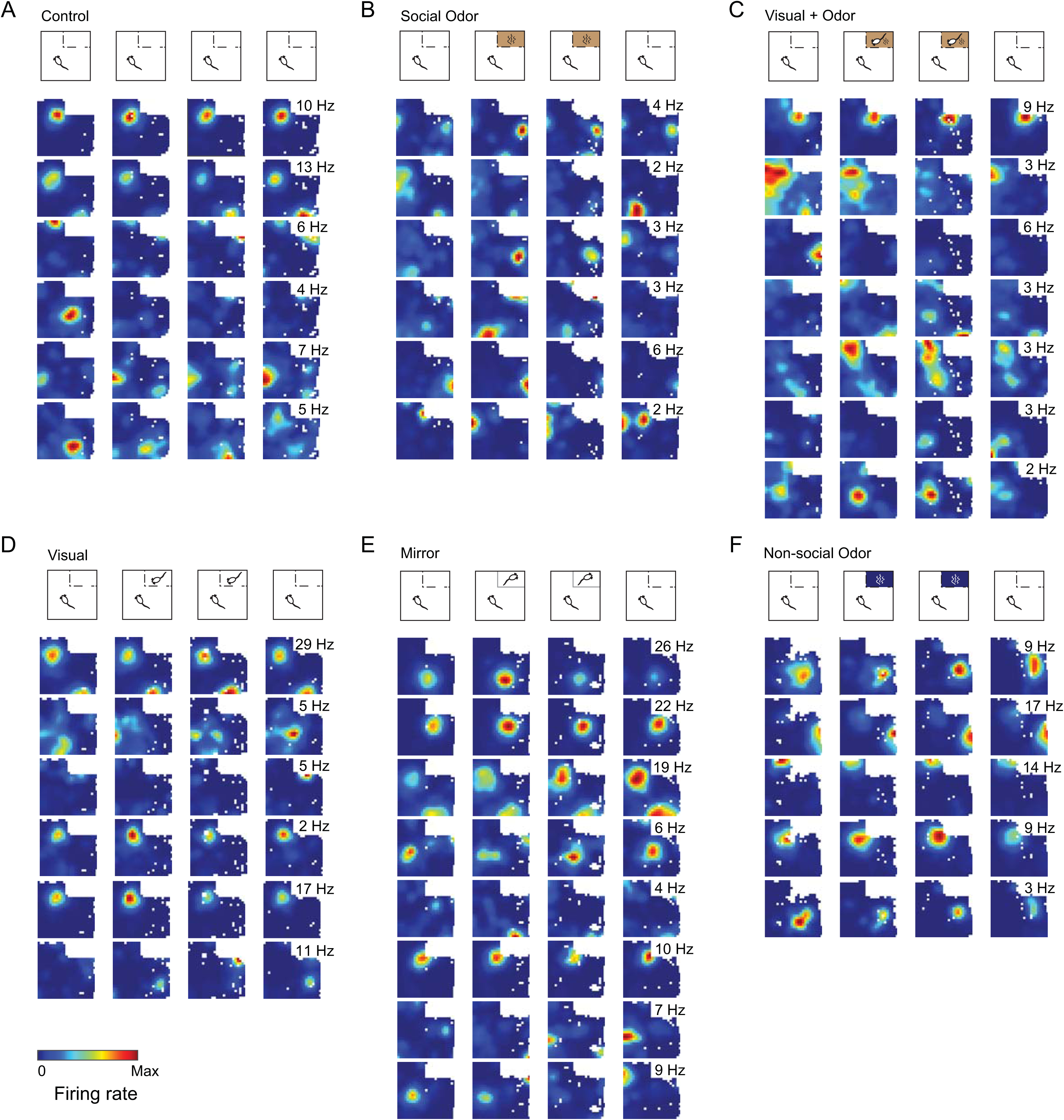
Example CA2 place cell firing rate maps across the four recording sessions for all conditions. Color-coded firing rate maps are shown for all place cells recorded on a single example tetrode in each condition for one example rat. Rate maps are shown scaled to the maximal firing rate (shown inset) of each cell across all sessions. White pixels indicate places that were not visited by the rat during a session.

**Figure 4.**
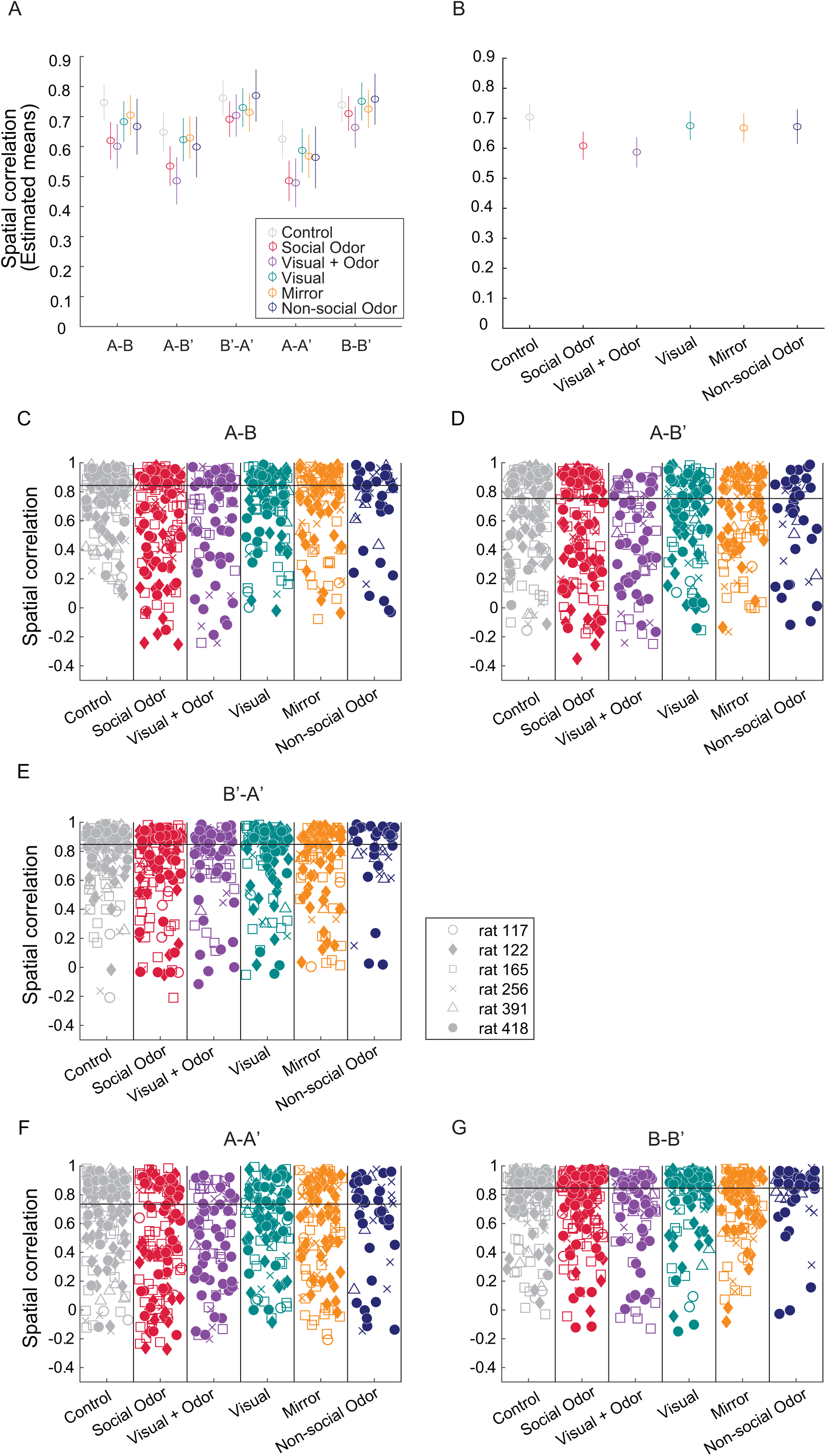
Remapping in CA2 in response to social stimuli. A-B. The estimated means of spatial correlation coefficients from our generalized linear mixed model are shown across each condition and session pair (A). The estimated means of spatial correlation coefficients from our generalized linear mixed model are shown across each condition for all session pairs combined (B). Error bars represent 95% confidence intervals. Statistics and associated p-values for specific pairwise comparisons of spatial correlations are shown in Table 3. C-G. Spatial correlation measures are shown for the entire sample of CA2 place cells for all pairs of sessions across all conditions. Each marker represents a spatial correlation value for an individual place cell. Different symbols are used for CA2 place cells recorded from different rats.

**Table 3.** Pairwise comparisons of spatial correlations between conditions.

| Pairwise contrast | t | df | Adjusted p-value |
| --- | --- | --- | --- |
| Control vs Social Odor | 6.3 | 3250 | 4.265E-9*** |
| Control vs Visual | 1.8 | 3250 | 0.411 |
| Control vs Visual + Odor | 6.4 | 3250 | 2.421E-9*** |
| Control vs Mirror | 2.3 | 3250 | 0.166 |
| Control vs Non-social Odor | 1.4 | 3250 | 0.770 |
| Social Odor vs Visual | -4.0 | 3250 | 0.001** |
| Social Odor vs Visual + Odor | 1.2 | 3250 | 0.935 |
| Social Odor vs Mirror | -3.6 | 3250 | 0.003** |
| Social Odor vs Non-social Odor | -2.8 | 3250 | 0.047* |
| Visual + Odor vs Visual | -4.5 | 3250 | 8.613E-5*** |
| Visual + Odor vs Mirror | -4.1 | 3250 | 0.001** |
| Visual + Odor vs Non-social Odor | -3.5 | 3250 | 0.004** |
| Visual vs Mirror | 0.4 | 3250 | 1.0 |
| Visual vs Non-social Odor | 0.1 | 3250 | 1.0 |
| Mirror – Non-social Odor | -0.1 | 3250 | 1.0 |

To quantify potential rate remapping across the various conditions, we computed a rate overlap measure across a pair of sessions for each cell. The rate overlap was defined as the ratio of the mean in-field firing rate of a place cell in each session in a session pair, with the smaller of the two firing rates in the numerator. To determine if rate overlap values differed across session pairs and conditions (i.e., Social Odor, Visual + Odor, Visual, Mirror, Non-social Odor, Control), we used a generalized linear mixed model statistical analysis (IBM SPSS Statistics, version 29.0.2.0). Condition and session pair were fixed factors, session pairs were repeated measures within rats, and multiple place cells were nested within rats. A condition by session pair interaction effect was also included in the model to determine whether differences between session pairs varied across conditions. The estimated mean rate overlap values for each condition and session pair from the generalized linear mixed model are shown in Supplementary Figure 2A. The estimated mean rate overlap values for each condition, collapsed across session pairs, are shown in Supplementary Figure 2B. Individual rate overlap values for each cell are shown for each session pair and each condition in Supplementary Figure 2C-G.

To determine if positions of place fields coherently shifted during remapping, we first identified cells that remapped and sorted them into “turn on”, “turn off”, and “field shift” categories (Supplementary Figure 3). Because place cells were classified as active if they reached a peak firing rate of at least 1 Hz, “turn on” cells were defined as cells that had a peak firing rate less than 1 Hz in Session A and greater than 1 Hz in Session B. We then calculated the distance between the position of the peak firing rate of the cell in Session B and the stimulus cage for “turn on” cells. “Turn off” cells were defined as cells that had a peak firing rate above 1 Hz in Session A and below 1 Hz in Session B. The distance between the position of the peak firing rate of the cell in Session A and the stimulus cage was calculated for “turn off” cells. Cells were identified as “field shift” cells if they reached a peak firing rate above 1 Hz in Sessions A and B but showed unstable place field locations. We assessed place field stability as follows, using criteria defined in a prior study (Widloski and Foster 2022). Cell IDs were randomly shuffled 1000 times within each condition and rat. Spatial correlation coefficients were then calculated for a given unit from session A and each shuffled rate map from session B. A place field was considered stable between sessions A and B if the spatial correlation coefficient exceeded the 95^th^ percentile of the shuffled distribution. For “field shift” cells, we calculated the distances between the stimulus cage and the positions of the peak firing rates of the cell in Sessions A and B. To determine if a cell’s place field moved closer to the stimulus cage when stimuli were presented, we calculated the difference between the distance estimates from Sessions A and B for “field shift” cells. We used the Shapiro-Wilk test of normality to determine if the distribution of distances was skewed within each condition (IBM SPSS Statistics, version 29.0.2.0). We additionally used an independent samples Kruskal-Wallis test to assess the extent to which distributions differed across conditions (IBM SPSS Statistics, version 29.0.2.0).

To quantify how selective a unit was for session A vs session B, we calculated a selectivity index for each unit, as in Hwaun and Colgin (2019). The selectivity index was defined as (μ_B_ – μ_A_)/(μ_B_ + μ_A_), where μ_A_ was the mean firing rate in session A and μ_B_ was the mean firing rate in session B. A value of −1 would indicate that a unit was exclusively active in session A, while a value of 1 would indicate that a unit was exclusively active in session B.

### 2.8. Place cell properties

Spatial information was calculated as previously described (Skaggs et al., 1996) using the following formula:

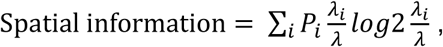

where *i* = the spatial bin index, *P_i_* = the probability of the rat being in the *i*th bin, *λ_i_* = the mean firing rate in the *i*th bin, and *λ* = the mean firing rate of the cell.

Place fields were separately identified for each cell in each session. Potential place fields were identified as contiguous bins of the smoothed rate map (see “*Place cell remapping analyses”* section) that exceeded 50% of the peak firing rate in that session. In order to be included for further analysis, the peak firing rate within a place field had to exceed 1 Hz, and the size of the place field had to exceed 10 cm^2^. To test if place field properties differed across conditions, we used a generalized linear mixed model statistical analysis (IBM SPSS Statistics, version 29.0.2.0). Multiple place cells were nested within rats, and sessions were repeated measures. A condition by session interaction effect was included in the model to determine whether differences between session pairs varied across conditions.

### 2.9. Exploration time

To investigate exploration of the stimulus cage, the arena was divided into 4 cm^2^ bins, as in a prior study (Zhu et al., 2023). The time spent in each bin during the first two minutes of every session was determined for each day. These exploration maps were then smoothed with a two-dimensional Gaussian kernel (standard deviation = 4 cm). Maps were averaged within and then across rats and plotted as a heat map for each condition (Figure 5A). The time spent within 12 cm of the cage was then calculated. A generalized linear mixed model analysis (IBM SPSS Statistics, V 29.0.2.0) was used to compare time spent investigating the stimulus cage across different sessions and conditions, with conditions and sessions as fixed factors. Conditions were repeated on different days within rats, with different days included as random factors within rats. Sessions were repeated measures within conditions. Post-hoc tests were performed to compare exploration times across conditions for the session of interest (i.e., Session B), using the Bonferroni correction for multiple comparisons.

**Figure 5.**
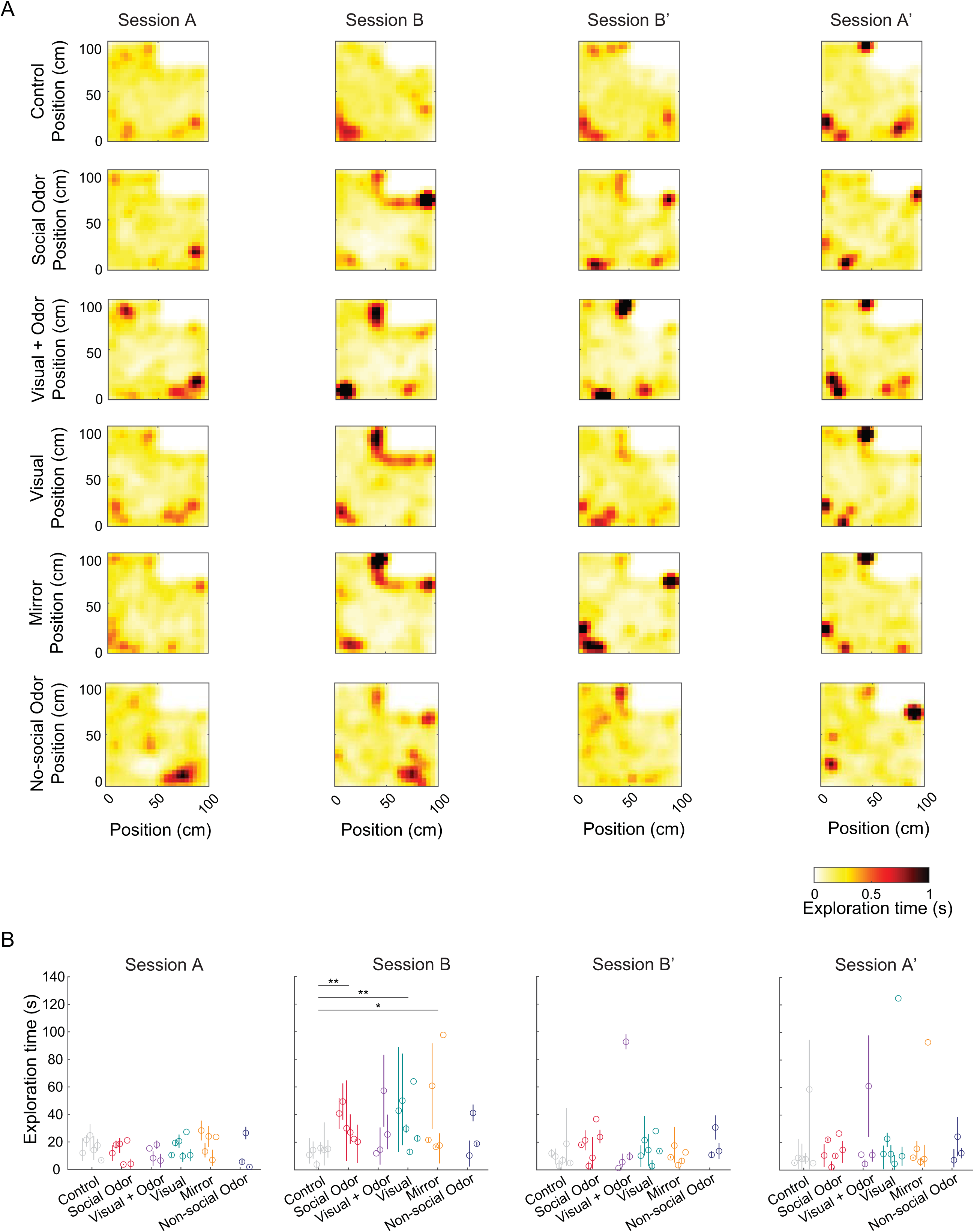
Exploration of stimuli across conditions. A. Heat maps show mean exploration times of different locations in the arena, with the stimulus cage shown in the top right corner of the arena. Time spent was calculated for the first 2 minutes of each session individually and then averaged within each rat. Heat maps shown are averaged across rats. B. Time spent exploring locations close to the cage (within 12 cm) was calculated for each session and then averaged within a rat. Individual dots represent the mean exploration time for each rat, and error bars represent 95% confidence intervals across all sessions within a rat. Rats increased their exploration time of the stimulus cage in session B for the Social Odor, Visual, and Mirror conditions compared to the Control condition (* indicates p-values < 0.05 and ** indicates p-values < 0.01).

### 2.10. Sharp wave-ripple detection and analysis

Sharp wave-ripples were detected from recordings obtained while rats rested in an elevated, towel-lined flowerpot outside of the arena. Detection and analysis methods were similar to previously published methods (Hwaun and Colgin, 2019). The LFP recorded on all tetrodes that had CA2 place cells was band-pass filtered between 150 and 250 Hz. A Hilbert transform was performed on the filtered LFP, and the absolute value of this signal was smoothed with a Gaussian kernel (standard deviation = 25 ms). Potential sharp wave-ripple events were detected when the signal exceeded at least 5 standard deviations above the mean and bounded by crossings of the mean. Overlapping events were combined across tetrodes, so the time interval of identified events could extend beyond sharp wave-ripples detected on a single tetrode (as in Hwaun and Colgin, 2019; Karlsson and Frank, 2009). Potential sharp wave-ripple events were kept for further analysis if they were between 50 and 500 ms in duration.

We then obtained the firing rate for each CA2 place cell around detected sharp wave-ripple events by constructing a spike raster with a bin size of 1 ms and smoothing the raster with a Gaussian kernel (standard deviation = 5 ms). To account for baseline firing rate differences among individual cells, the firing rate was normalized by the average firing rate 400-100 ms before ripple onset (Hwaun and Colgin, 2019). The peak normalized firing rate was obtained for each unit by taking the maximum normalized firing rate after sharp wave-ripple onset. Because a broad range of peak normalized firing rates was observed, we rescaled the distribution using a log_10_ transformation of the peak normalized firing rates. To determine the extent to which sharp wave-ripple-associated firing rates of individual cells were altered by presentation of a social stimulus, we computed the difference between the peak normalized firing rates during the rest periods after sessions A and B (i.e., peak normalized firing rate after session B - peak normalized firing rate after session A) for the Control and Social Odor conditions. A positive value indicates that a cell had higher sharp wave-ripple-associated firing in the rest session following session B, and a negative value indicates that a cell had higher sharp wave-ripple-associated firing in the rest session following session A. We used multiple linear regression (IBM SPSS Statistics, V29.0.2.0) to assess the effects of session selectivity (as measured using the selectivity index, see *Place cell remapping analyses* section) and experimental condition on the difference in sharp wave-ripple-associated firing between rest sessions (as in Hwaun and Colgin, 2019). The regression analysis was only performed for conditions that had more than 100 active cells during both run and rest sessions (Control condition: n = 130 cells from 5 rats, Social Odor condition: n = 132 cells from 5 rats).

### 2.11. Data and Code Availability

MATLAB (Mathworks) scripts were custom written for the analyses in this study, based on algorithms that have been used in prior studies, as described above. Scripts and data are available upon request.

## 3. Results

In our prior study of CA2 place cells (Alexander et al., 2016), we compared CA2 place cell responses to a rat in a soiled home cage and responses to the presentation of an object that resembled a rat, namely a stuffed toy rat. However, the stuffed toy lacked some visual components that occur during interactions with a live rat (e.g., motion content). Also, the toy rat was presented in a clean cage and thus lacked the olfactory components of a social experience. The major goal of the present study was to determine the extent to which different sensory modalities associated with a social experience drive CA2 place cell remapping to social stimuli. We employed an open field exploration behavior paradigm (see “*Behavioral task“* section of Materials and Methods) in which different types of stimuli were presented across sessions for various experimental conditions (i.e., Social Odor, Visual + Odor, Visual, Mirror, Non-social Odor) and compared to a Control condition in which no stimuli were presented (Figure 1).

### 3.1. A significant proportion of CA2 place cells globally remapped to social odor stimuli

The olfactory system is highly important for social recognition and social behaviors in rodents (Oettl and Kelsch, 2018). Therefore, we hypothesized that olfactory cues would be a key sensory component of CA2 place cell responses to social stimuli. In the Social Odor condition, rats explored an environment in which social odors were presented, but another rat was not present. This condition isolated the olfactory content of a social interaction and preserved the ethological relevance of the presented stimuli. Example CA2 place cell firing rate maps during presentation of social odors (Figure 3B) show that a subset of cells changed the location of their firing fields, or globally remapped, when social odors were presented. To quantify global remapping in CA2 place cells, we compared firing rate map changes in cells recorded in the Social Odor condition (145 cells in 6 rats, Table 1) to firing rate map changes in cells recorded in the other conditions (see Figure 3 for example rate maps for all conditions and Table 1 for cell yields). We calculated the spatial correlation between rate maps from pairs of sessions for each cell (Figure 4). We found that spatial correlation values for the Social Odor condition were significantly lower than the Control, Visual, Mirror, and Non-social Odor conditions (generalized linear mixed model, no significant interaction between session pair and condition, F(20, 3250) = 0.843, p = 0.662 (Figure 4A); significant main effect of condition, F(5, 3250) = 12.7, p < 0.001 (Figure 4B); significant differences in post-hoc tests for Social Odor vs. Control (t(3250) = 6.3, p < 0.001), Social Odor vs. Visual (t(3250) = 4.0, p = 0.001), Social Odor vs. Mirror (t(3250) = 3.6, p = 0.003), and Social Odor vs. Non-social Odor (t(3250) = 2.8, p = 0.047)). In agreement with our prior findings, CA2 place cells also showed significant global remapping when a familiar conspecific rat was presented in a soiled home cage containing social odors (Visual + Odor condition, Figures 3C and 4, significant differences in post-hoc tests for Visual + Odor vs. Control (t(3250) = 6.4, p < 0.001), Visual + Odor vs. Visual (t(3250) = 4.5, p < 0.001), Visual + Odor vs. Mirror (t(3250) = 4.1, p = 0.001), and Visual + Odor vs. Non-social Odor (t(3250) = 3.5, p = 0.004)). In contrast, significant global remapping was not observed when a stimulus rat was presented in a clean cage with a filter top, a condition that maintained all visual components of social interactions and minimized social odors (Visual condition, Figures 3D and 4, no significant differences in post-hoc test for Visual vs. Control (t(3250) = 1.8, p = 0.411)). Similarly, CA2 place cell firing rate maps did not change significantly when a stimulus rat was presented in a clean cage with a filter top and a one-way mirror to prevent visually mediated reciprocal interactions between rats (Mirror condition, Figures 3E and 4, no significant differences in post-hoc test for Mirror vs. Control (t(3250) = 2.3, p = 0.166)). To test whether remapping of CA2 place cell responses was specific to social odors, we presented a non-social odor in the stimulus cage. We found that presentation of a non-social odor did not significantly change CA2 place cell firing rate maps (Non-social Odor condition, Figures 3F and 4, no significant differences in post-hoc test for Non-social Odor vs. Control (t(3250) = 1.4, p = 0.770)) (see Table 3 for all pairwise comparisons). Together, these results show that a subset of CA2 place cells globally remapped when social odors were presented but not during interactions with another rat when social odors were minimized.

While non-odorant social conditions and non-social odors were not found to induce global remapping in CA2 place cells, it remains possible that these conditions induced rate remapping in CA2. To test this possibility, we compared rate overlap values across conditions (Supplementary Figure 2). Consistent with previous work (Alexander et al., 2016), we found that none of the social conditions induced significant rate remapping in CA2 place cells. Further, the presentation of a non-social odor did not induce significant rate remapping in CA2 place cells (generalized linear mixed model, no significant interaction between session pair and condition, F(20,3213) = 0.7, p = 0.803; no main effect of condition, F(5,3213) = 2.0, p = 0.076)).

### 3.2. No coherent movement of place fields during remapping

CA1 place fields have been reported to shift with salient stimuli (Fenton et al., 2000; O’Keefe and Conway, 1978) or move toward locations of salient rewarding stimuli (Breese et al., 1989; Fenton et al., 2000). However, our previous work showed that fields of CA2 place cells in rats did not move closer to a social stimulus during social remapping (Alexander et al., 2016). Conversely, recordings of CA2 place cells in mice showed that a subpopulation of CA2 place cells remapped when a conspecific mouse was moved to a new location of the arena, and this subpopulation of “social-remapping” cells had fields significantly closer to the stimulus mouse than fields of non-remapping (“social-invariant”) cells (Oliva et al. 2020). To test if CA2 place cells moved coherently towards or away from the stimulus cage in our experiment, we first identified place cells that remapped between sessions A and B. These cells were identified as cells that either gained a place field (“turn on” cells), lost their place field (“turn off” cells), or shifted their place field (“field shift” cells). For “turn on” cells, we calculated the distance between the cell’s peak firing rate position in the session in which stimuli were presented (Session B) and the position of the stimulus cage (Supplementary Figure 3A). Similarly, we calculated the distance between the peak firing rate position and the position of the stimulus cage in Session A for the “turn off” cells (Supplementary Figure 3B). If place cells preferentially gained or lost fields close to the stimulus cage, we would have expected skewed distributions towards low values, which we did not observe. However, definitive conclusions cannot be drawn from this observation given the low number of cells that met the criteria for classification as “turn on” or “turn off” cells.

To determine whether CA2 place cells preferentially shifted their place fields closer to the stimulus cage, we estimated the distance between the stimulus cage and cells’ peak firing rate positions in Sessions A and B and then calculated the difference between these two distance estimates (Supplementary Figure 3C). If place fields shifted closer to the cage, we would have expected values to skew negatively. However, differences between place field distances from the cage between Sessions A and B were normally distributed for all conditions (Shapiro-Wilk test of normality, Control: W(20) = 1.0, p = 0.717; Social Odor: W(30) = 1.0, p = 0.339; Visual + Odor: W(16) = 0.9, p = 0.135; Visual: W(15) = 0.9, p = 0.142; Mirror: W(18) = 0.9, p = 0.346; Non-social Odor: W(4) = 0.880; p = 0.340). Further, the distributions of distance shifts did not differ significantly across conditions (Kruskal-Wallis test, H(5) = 6.3, p = 0.278). Therefore, consistent with our previous findings (Alexander et al., 2016), no coherent movement of place fields toward or away from the stimulus cage was observed. However, our experimental design did not allow us to identify “social-remapping” and “social-invariant” place cells as was done in a previous study (Oliva et al. 2020). That is, social-remapping cells in the previous study in mice were defined as place cells that displayed place fields that shifted in a manner that preserved a fixed distance from a stimulus mouse, whereas place fields of “social-invariant” cells maintained a fixed location in the environment. In the present experiment, only one stimulus was presented in each session, and the location where it was presented did not change across sessions. It is possible that the present sample of cells contained a mixture of social-remapping and social-invariant place cells and that potential shifts in field relative to the social stimulus were masked as a result. Alternatively, the lack of a preferential shift in place fields near the social stimulus may be due to the nature of odor stimuli, as airborne odorants diffuse across widespread locations within an environment. This may allow a rat to perceive odors regardless of their exact location in the arena. It is also possible that the introduction of social odors to an environment was sufficient to change its context as a whole. This generalizability may help rats to associate conspecifics with larger areas of an environment.

### 3.3. CA2 place cell properties were not modulated by social stimuli

In addition to remapping, we examined general CA2 place cell properties during presentation of social and non-social stimuli. We found that spatial information was unaffected by the inclusion of social stimuli or a non-social odor in an environment (Supplementary Figure 4A; generalized linear mixed model, no significant interaction between session and condition (F(23,2596) = 1.0, p = 0.369)). We also saw no significant changes in peak firing rates (Supplementary Figure 4B; generalized linear mixed model, no significant interaction between session and condition (F(23,2413) = 1.0, p = 0.478)), mean in-field firing rates (Supplementary Figure 4C; generalized linear mixed model, no significant interaction between session and condition (F(23,2413) = 1.0, p = 0.477)), or the number of place fields per cell (Supplementary Figure 4D; generalized linear mixed model, no significant interaction between session and condition (F(23, 2596) = 0.9, p = 0.583)). These results suggest that CA2 place cells represented spatial locations similarly in social and non-social conditions.

### 3.4. Rats explored all modalities of social stimuli

It is possible that the lack of remapping to non-olfactory social stimuli may be due to inadequate exploration of these stimuli. To test this possibility, we examined rats’ stimuli exploration time during the first 2 minutes of each exploration session for each experimental condition (Figure 5). We found that stimulus cage exploration time during session B of most social conditions (i.e., Social Odor, Visual, and Mirror) significantly increased compared to session B of the Control condition (generalized linear mixed model, interaction between session and condition (F(15,284) = 2.3, p = 0.004); post-hoc tests, session B: significant difference between Control vs. Social Odor (t(284) = 3.0, p = 0.037); Control vs. Visual (t(284) = 3.5, p = 0.009); and Control vs. Mirror (t(284) = 3.4, p = 0.010); no significant difference between Control vs. Visual + Odor (t(284) = 2.2, p = 0.325); no significant difference between Control vs. Non-social Odor (t(284) = 1.2, p > 0.999)). It is important to note that stimulus cage exploration times significantly increased in the social conditions in which CA2 place cells did not show significant remapping (i.e., Visual and Mirror conditions), suggesting that the lack of place cell remapping in these conditions was not due to reduced exploration of social stimuli. Note also that exploration times did not significantly differ between Visual + Odor and Control conditions, although significant remapping was observed in the Visual + Odor condition. The insignificant difference in exploration times may be due to the lower number of rats in the Visual + Odor condition (i.e., 4 rats in Visual + Odor condition compared to 5-6 rats in other social conditions, see Table 1). Similarly, exploration times also did not differ between Non-social Odor and Control conditions, perhaps due to the lower number of rats in the Non-social Odor condition (i.e., only 3 rats, see Table 1). Alternatively, the lack of ethological relevance of the hexyl acetate odor may have contributed to a lack of increased exploration of the stimulus cage during presentation of the Non-social Odor.

### 3.5. Place cells that responded to social odors were preferentially recruited into sharp wave-ripples

CA2 place cells that encode a social experience with a novel conspecific have been shown to reactivate during sharp wave-ripples in mice (Oliva et al., 2020). Therefore, we aimed to determine whether presentation of social odors alone was sufficient to drive preferential sharp wave-ripple-associated reactivation of CA2 place cells that responded to social odors. We used a selectivity index (see Materials and Methods) to classify cells according to their firing preferences during the Control and Social Odor conditions. The selectivity index defined the extent to which CA2 place cells fired during session A (i.e., the session in which an empty cage was presented) compared to session B (i.e., the session in which a cage containing social odors was presented in the Social Odor condition). We then estimated the normalized average peak firing rate of each cell during sharp wave-ripples in rest sessions after session A and session B (see Materials and Methods). Next, we used the difference between a cell’s peak normalized firing rate during the rest session after session B and the rest session after session A as a measure of the extent to which a cell changed its sharp wave-ripple-associated firing after presentation of social odors. CA2 place cells show unstable firing patterns in unchanged environments over time (Mankin et al., 2015). Thus, we performed the same analysis for CA2 place cells recorded in the Control condition to ensure that changes in sharp wave-ripple-associated firing were not explained by the effect of time. We found that a cell’s selectivity for social stimuli in active exploratory sessions correlated with sharp wave-ripple-associated peak firing rate changes in the Social Odor condition but not the Control condition (Figure 6; multiple linear regression, F(3,243) = 2.661, p = 0.049; interaction between selectivity index and condition, t = 2.448, p = 0.015, Pearson correlation between selectivity index and firing rate changes for the Control condition R = −0.102, p = 0.266, Pearson correlation between selectivity index and firing rate changes for the Social Odor condition R = 0.227, p = 0.010). Specifically, cells that increased their firing rates during exploration of a social odor preferentially increased their sharp wave-ripple-associated firing rates during rest following social exploration. These results suggest that the presentation of a social odor alone is sufficient to preferentially reactivate CA2 place cells that code social stimuli.

**Figure 6.**
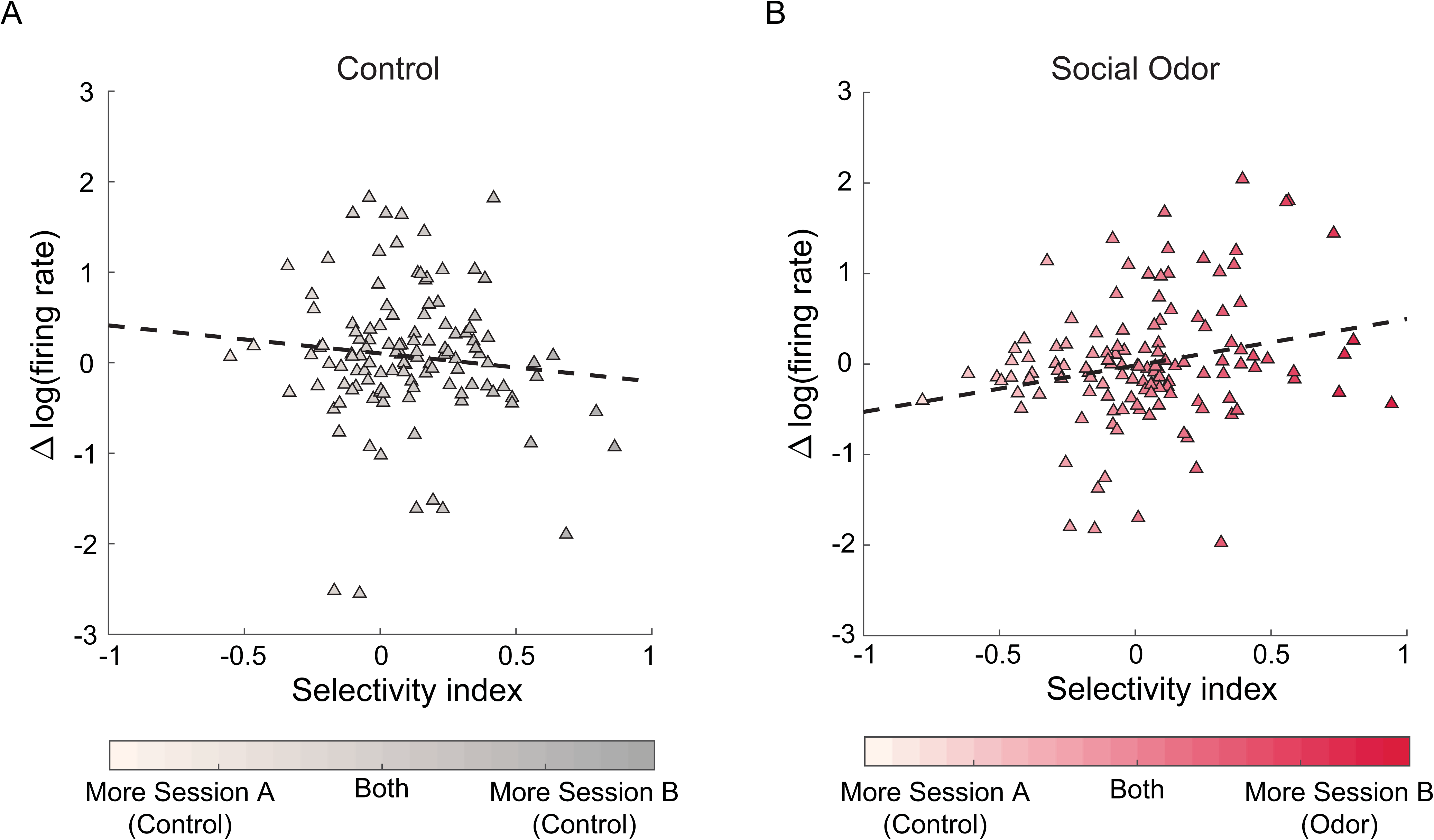
CA2 place cells that responded to a social odor preferentially fired during sharp wave-ripples. A. The difference between firing rates of CA2 place cells during sharp wave-ripples in the rest periods following sessions A and B were positively correlated to the selectivity index for the Social Odor condition (B) but not the Control condition (A). The selectivity index indicates a cell’s preference for session A (value of −1 for maximum selectivity in session A) or session B (value of 1 for maximum selectivity in session B).

## 4. Discussion

An increasing number of lesion, genetic, optogenetic, and pharmacogenetic manipulation studies have implicated CA2 in social memory processing (Hitti and Siegelbaum, 2014; Meira et al., 2018; Oliva et al., 2020; Smith et al., 2016; Stevenson and Caldwell, 2014). In this study, we expanded on prior results showing that CA2 place cells remap during a social interaction (Alexander et al., 2016) by investigating how various sensory modalities of social stimuli contribute to this response. Here, we show that CA2 place cell remapping is primarily driven by the olfactory component of social stimuli. Exposure to social odors in the absence of a rat was sufficient to induce remapping in CA2 place cells. CA2 place cells did not respond to social interactions with another rat when social odors were reduced. The lack of remapping when social odors were reduced was not explained by inadequate exploration of social stimuli. Lastly, CA2 place cells that showed increased firing during exploration of social odors were preferentially reactivated during sharp wave-ripples in subsequent rest. Our findings support the notion that CA2 place cells may integrate social olfactory information with contextual information to encode and consolidate social episodic memories (Oliva, 2022; Oliva et al., 2023). However, future experiments that selectively silence CA2 are needed to support a direct link between CA2 place cell activity and behavioral performance on memory tasks requiring association of social odors with specific contexts. Also, while social experiences did not induce significant global remapping in CA2 place cells when social odors were lacking, the possibility remains that other sensory modalities of a social experience are processed via different pathways and brain structures and thereby contribute to social recognition and social memory processes.

The sensory modalities that drive social recognition vary across species. While humans depend predominantly on auditory and visual cues for social recognition, rodents and other mammals rely heavily on chemosensory cues in the form of olfactory or pheromonal signals (Popik and van Ree, 1998). Indeed, olfactory bulb lesions or chemically induced anosmia impair individual recognition in rats (Dantzer et al., 1990; Popik et al., 1991). The complex mix of chemosensory signals embedded in rat urine convey information about a rat’s social status and identity (Hurst, 2005). CA2 neurons differentially respond to urine from different conspecifics in mice (Hassan et al., 2023). The ability of CA2 to respond to social odors in the absence of conspecifics, as seen in our study, may allow rats to discern territory and identity information, which may drive social behavior even when other rats are not present. The precise mechanisms underlying social odor detection in CA2 neurons are unknown but may be related to the distinctive neurochemical and structural features of CA2 neurons.

CA2 neurons contain an abundance of neuropeptide receptors, including oxytocin receptors (OXTR) and vasopressin receptors (AVP1-b) (Mitre et al., 2016; Vaccari et al., 1998; Young and Song, 2020). Genetic or pharmacological ablation of CA2 OXTRs impairs the persistence of long-term social recognition memory and the ability to discriminate between social stimuli in mice (Lee et al., 2008; Lin et al., 2018; Raam et al., 2017). AVP1-b activation in CA2 has been associated with social aggression (Pagani et al., 2015) and the enhancement of social memory (Smith et al., 2016). Long-range axonal projections from the paraventricular nucleus of the hypothalamus (PVN) may be the source of neuropeptide release onto CA2 neurons during social interactions. The PVN receives input from the olfactory system (Guevara-Aguilar et al., 1988) and directly innervates CA2 (Cui et al., 2013; Zhang and Hernández, 2013).

Several other brain regions that are involved in social processing send direct inputs to CA2, including the hypothalamic supramammillary nucleus (SuM) and the lateral entorhinal cortex (LEC). SuM projections to CA2 are preferentially activated by novel social encounters (Chen et al., 2020), making SuM inputs an unlikely candidate to induce CA2 place cell responses to a familiar social odor. Conversely, neuronal activity in the LEC showed similar increases during social exploration for both novel and familiar conspecifics (Lopez-Rojas et al., 2022). Recent studies have suggested that the direct projection from LEC to CA2 is essential for social memory (Dang et al., 2022; Lopez-Rojas et al., 2022). Whether LEC input to CA2 is selectively enhanced during exploration of a social odor alone remains to be tested, but a more general involvement of the LEC in odor processing is well documented (Igarashi et al., 2012; Kerr et al., 2007).

CA2 also receives input from granule cells of the dentate gyrus (DG). Both mature and adult-born granule cells directly synapse onto CA2 neurons via mossy fiber projections (Kohara et al., 2014; Llorens-Martín et al., 2015; Laham et al., 2024). The DG is important for associating odors with larger contexts in discrimination tests (Morris et al. 2012). In addition, recent work has shown that adult-born granule cell projections to CA2 are necessary for the retrieval of social memories formed during development (Laham et al., 2024). The exact relationship between the DG and CA2 in social odor processing is currently unknown and warrants further investigation.

Consistent with previous work, our results show that CA2 place cells that remap in response to a familiar social odor do not return to their original activity, despite the removal of the social odor in the last session (Alexander et al., 2016). It is possible that the persistent change in CA2 activity is a result of lingering social odors in the arena. Recording CA2 place cells for longer than 20 minutes after the removal of the odor would address if CA2 place cell activity eventually returns to baseline, although this would be complicated by the effect of time on CA2 place cells (Mankin et al., 2015). However, this persistent change in CA2 place cell activity may also suggest the involvement of synaptic plasticity. Interestingly, CA2 synapses are resistant to standard long-term potentiation (LTP) compared to other hippocampal subfields (Chang et al., 2007; Zhao et al., 2007). How, then, are long-lasting changes occurring in CA2 place cells? One hypothesis is that the release of social neuropeptides, such as vasopressin and oxytocin, can promote potentiation of CA2 neurons (Lin et al., 2018; Pagani et al., 2015; Tirko et al., 2018). Selective activation of OXTRs or AVP1-b *in vitro* robustly depolarizes CA2 pyramidal neurons and lowers the threshold for LTP at excitatory synapses (Dang et al., 2022; Tirko et al., 2018). As a result, neuropeptide receptor activation may refine the responsiveness of CA2 pyramidal neurons to synaptic input from upstream structures, thus creating appropriate conditions for synaptic plasticity.

A question remains of how social information coded by CA2 is transmitted to downstream regions that control behavior. The main output of dorsal CA2 is dorsal CA1. However, at the single cell level, dorsal CA1 place cell activity was unaffected by social interactions (Alexander et al., 2016). Dorsal CA2 neurons also project to ventral CA1, a region that is essential for social memory (Okuyama et al., 2016). Recent work has shown that ventral CA1 place cells did not remap to a social stimulus (Wu et al, 2023). However, ventral CA1 place cells became significantly more spatially selective in an environment that contained a social stimulus compared to an empty environment (Wu et al. 2023). Further, the activity of a subset of ventral CA1 neurons was modulated by the presence of a conspecific (Rao et al. 2019; Wu et al. 2023). Inhibition of dorsal CA2 projections to ventral CA1 impairs social memory (Meira et al., 2018; Tsai et al., 2022), raising the possibility that social remapping in dorsal CA2 induces the alterations in neuronal firing in ventral CA1 in response to social stimuli.

Interestingly, recent work has shown that sharp wave-ripples that originate in dorsal CA2 are important for social memory consolidation (Oliva et al., 2016, 2020). While a portion of the sharp wave-ripples initiated in dorsal CA2 propagate to other subregions of the dorsal hippocampus, some do not propagate to dorsal CA1 or dorsal CA3 (Oliva et al., 2016). Given that individual place cells in dorsal CA1 did not remap to a social stimulus (Alexander et al., 2016), it is possible that the reactivation of CA2 place cells that respond to a social odor may be limited to sharp wave-ripples that do not propagate to dorsal CA1. Instead, the function of these sharp wave-ripples may be to propagate information to ventral CA1 in order to consolidate memories of a social experience (Meira et al., 2018). Consistent with this idea, the rate of sharp wave-ripples increased in ventral CA1 when conspecifics were present (Rao et al., 2019). A future study examining the reactivation of dorsal CA2 place cells that respond to social stimuli while simultaneously recording place cells in dorsal and ventral CA1 will be essential for our understanding of how reactivation of CA2 place cells during sharp wave-ripples supports social memory consolidation.

While hippocampal place cells are able to represent position and respond to social stimuli such as odors at the single cell level, as evidenced by place cell remapping, population-level activity in the hippocampus may incorporate the activity of both place cells and non-spatially modulated cells to encode other features of an environment and ongoing behavior (Meshulam et al., 2017; Nagelhaus et al., 2023; Stefanini et al., 2020). The relatively low CA2 cell yields per rat in our data (see Table 1) made it difficult to examine population-level activity in CA2 across conditions. While we did not see remapping at the single cell level for our social conditions that minimized social odors and for our non-social odor condition, the possibility remains that these conditions induced population-level changes in CA2 activity.

Another important consideration for this study is that rats were singly housed after surgery to reduce trauma to the surgical site and protect drive implants. Social isolation can impair the persistence of social recognition memory in both mice and rats (Kogan et al. 2000; Shahar-Gold et al. 2013). This impairment may be due to a de-coupling of the olfactory bulb and dorsal hippocampus in mice (Almeida-Santos et al. 2019). We attempted to minimize the effects of social isolation by housing familiar conspecifics in adjacent cages, but we cannot rule out unintended consequences of social isolation on CA2 responses to social stimuli.

Another important question left unanswered is the extent to which CA2 is specialized for social memory. The present results show that a social odor, but not a non-social odor, was sufficient to induce CA2 place cell remapping. However, due to the differences in ethological relevance between social and non-social odors in this study, it remains unclear whether non-social odors of comparable salience or ethological relevance would affect CA2 place cell activity. Future studies examining the activity of CA2 place cells in response to ethologically relevant non-social odors, such as predator urine, will help determine whether CA2 integrates other salient odors into representations of space or whether the role of CA2 is specific to social information.

The current study shows how presentation of a social odor alone provides a well-controlled social stimulus to induce changes in place cell activity in dorsal CA2. Use of a social odor can eliminate confounds that could be introduced by the presence of a conspecific, such as variations in locomotor behavior, social interactions, and ultrasonic vocalizations. Presentation of a social odor thereby provides a valuable paradigm for assessing CA2 physiology in rodent models of diseases and disorders, such as autism spectrum disorders, involving aberrant processing of social stimuli.

## CRediT authorship contribution statement

**Emma Robson**: Formal analysis, investigation, writing – original draft, visualization. **Margaret M. Donahue:** Formal analysis, investigation, writing – original draft, visualization, funding acquisition. **Alexandra J. Mably:** Conceptualization, methodology, formal analysis, investigation. **Peyton G. Demetrovich:** Investigation. **Lauren T. Hewitt:** Investigation. **Laura Lee Colgin:** Conceptualization, methodology, resources, writing – review and editing, supervision, project administration, funding acquisition.

## Acknowledgements

The authors thank Isabella Lee, Misty Hill, Ayomide Akinsooto, Sirisha Dhavala, and Alexa Hassien for technical assistance with this project and Chenguang Zheng and Ernie Hwaun for providing MATLAB code for some of the analyses. The authors acknowledge the Texas Advanced Computing Center (TACC) at The University of Texas at Austin for providing data storage resources that have contributed to the research described within this article. URL: http://www.tacc.utexas.edu

## Funding

This research was supported by the Department of Defense (grant number W81XWH1810314) and the National Institutes of Health (grant numbers R56MH125655, R01MH131317, and F31MH127933).

## Data availability

Data will be made available on request.

---

**Supplementary Figure 1.**
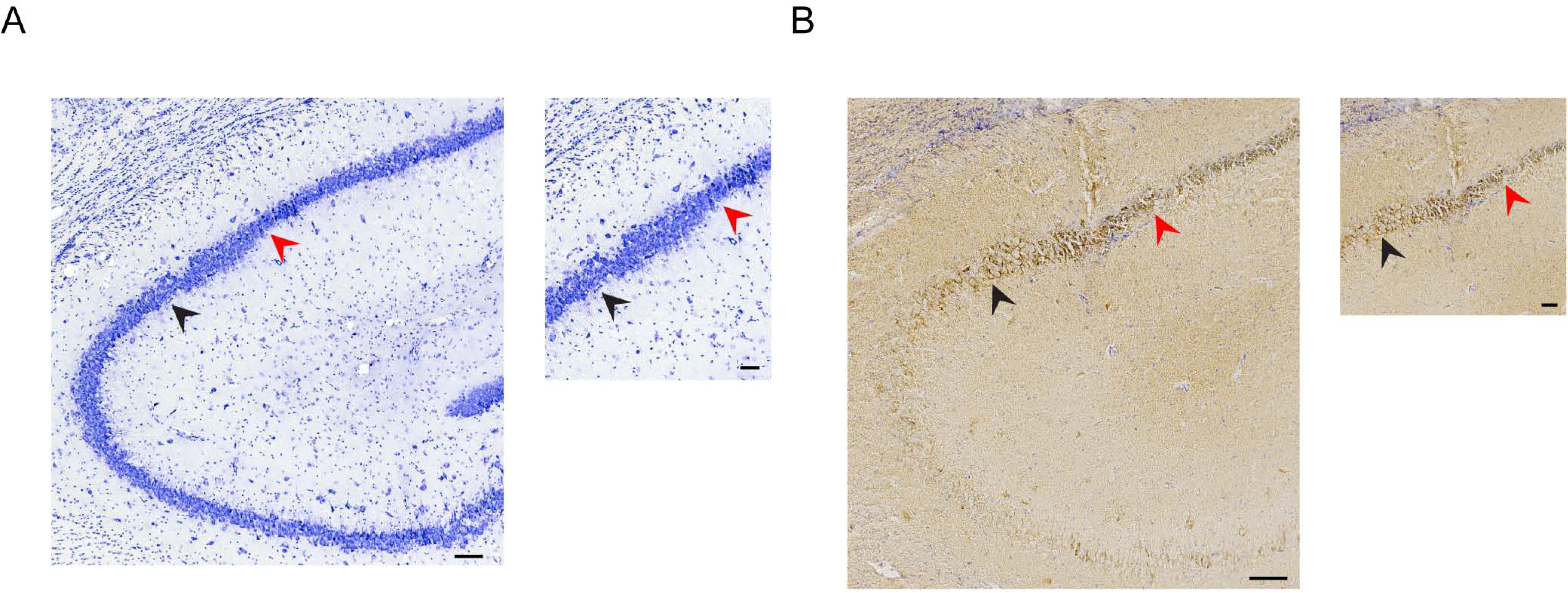
Example of CA2 localization using the CA2 marker STEP. A. A Cresyl violet-stained brain section from an example rat (rat 117) showing CA3/CA2 (black arrows) and CA2/CA1 (red arrows) borders. Scale bar, 100 µm left and 50 µm right. B. Example tetrode track in CA2 from rat 117. Sections were immunostained with a CA2 marker, STEP, and developed with 3,3-diaminobenzidine (brown). Sections were counter-stained with cresyl-violet (purple/blue). Scale bar, 100 µm left and 50 µm right.

**Supplementary Figure 2.**
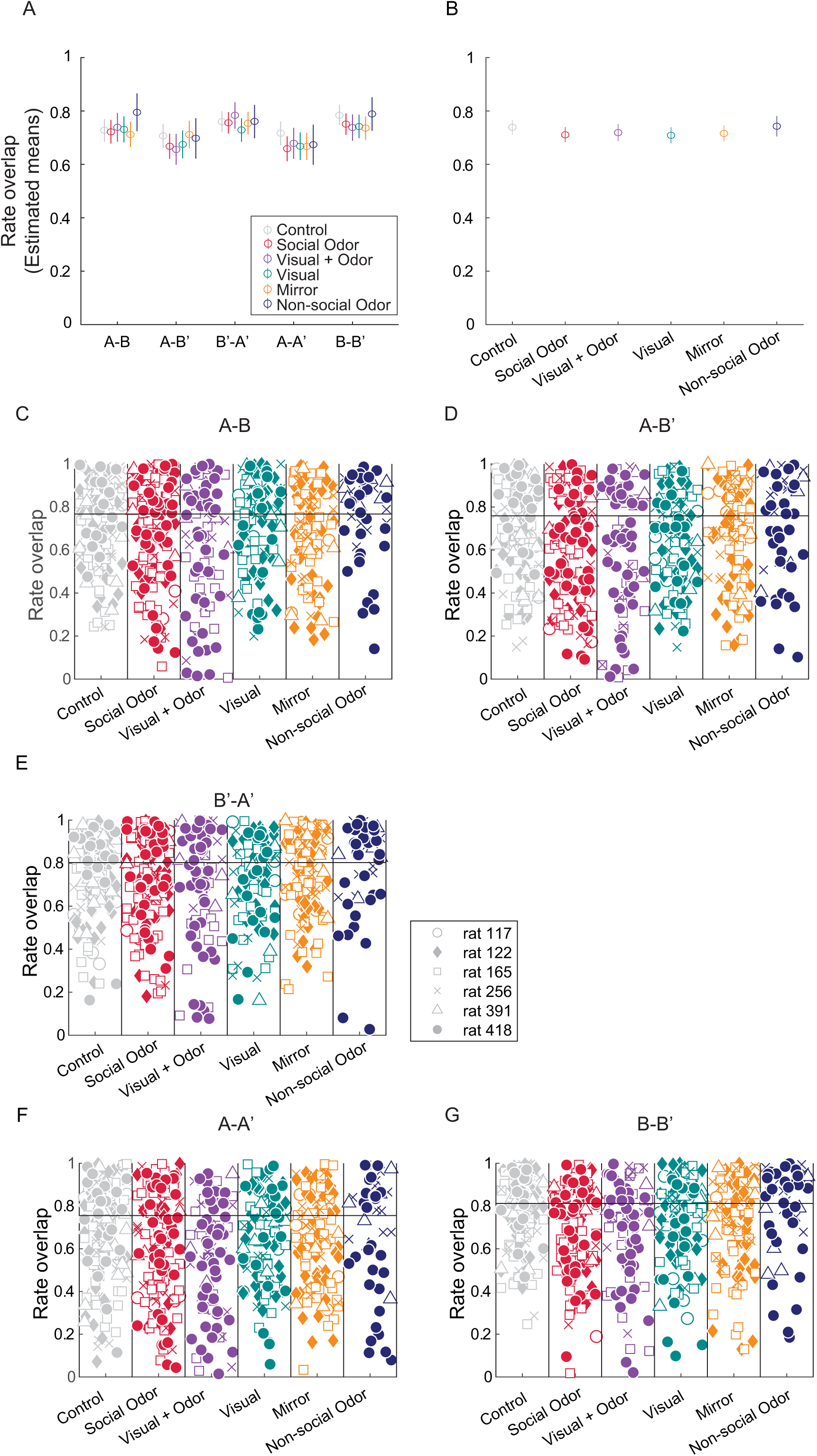
No rate remapping was observed in CA2 in response to social stimuli. A-B. The estimated means of rate overlap values from a generalized linear mixed model are shown across each condition and session pair (A). The estimated means of rate overlap values from the generalized linear mixed model are shown across each condition for all session pairs combined (B). Error bars represent 95% confidence intervals. C-G. Rate overlap values are shown for the entire sample of CA2 place cells for all pairs of sessions across all conditions. Each marker represents a rate overlap value for an individual place cell. Different symbols are used for CA2 place cells recorded from different rats.

**Supplementary Figure 3.**
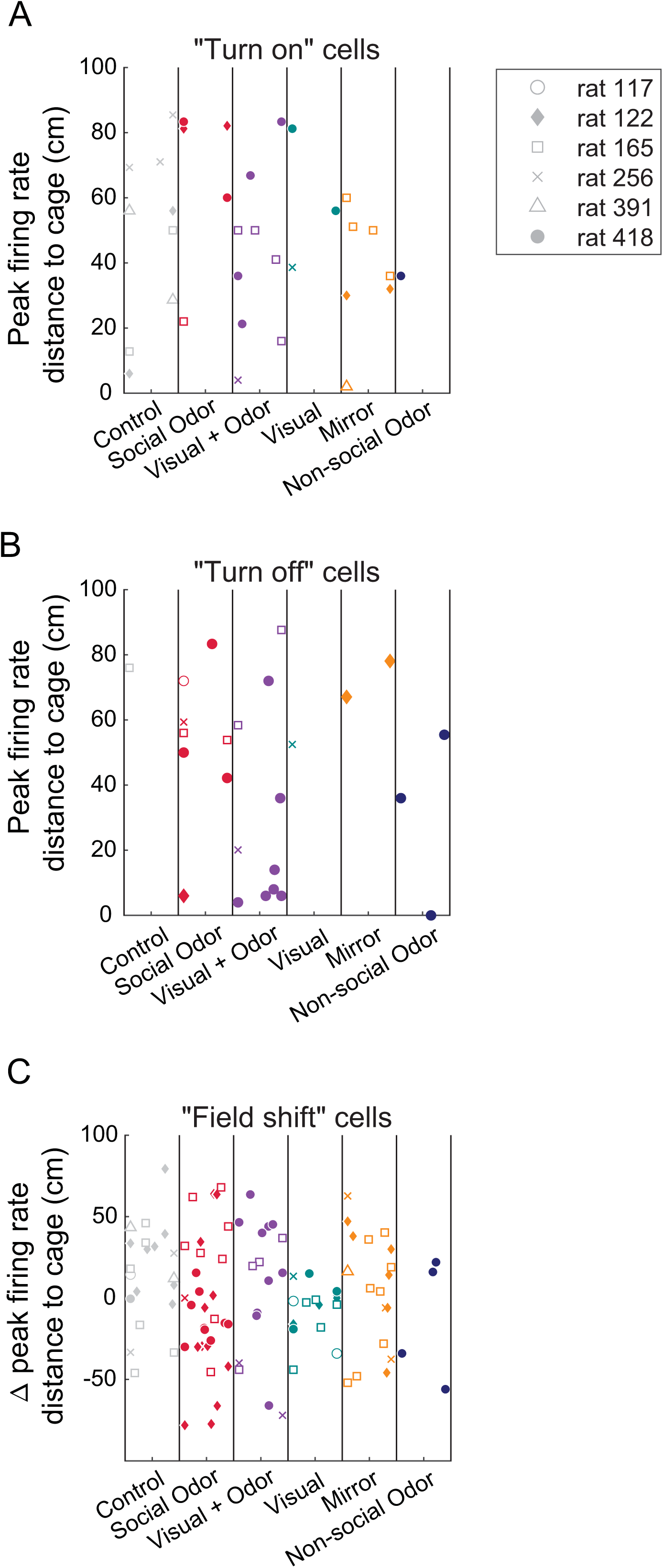
Evaluation of CA2 place field shifts relative to the location of the stimulus cage. A. Shown are the distances measured between the positions of peak firing in Session B and the stimulus cage for CA2 place cells that “turn on”, or gain a field, in Session B. B. Shown are the distances between the peak firing rate position in Session A and the location of the stimulus cage for CA2 place cells that “turn off”, or lose a field, in Session B. C. Shown are the changes in distance from the stimulus cage of the peak firing rate positions in Sessions A and B for CA2 place cells that were active in both Sessions A and B but shifted their place field locations. Individual markers represent measurements from individual place cells. Different symbols represent distance measurements for CA2 place cells recorded from different rats.

**Supplementary Figure 4.**
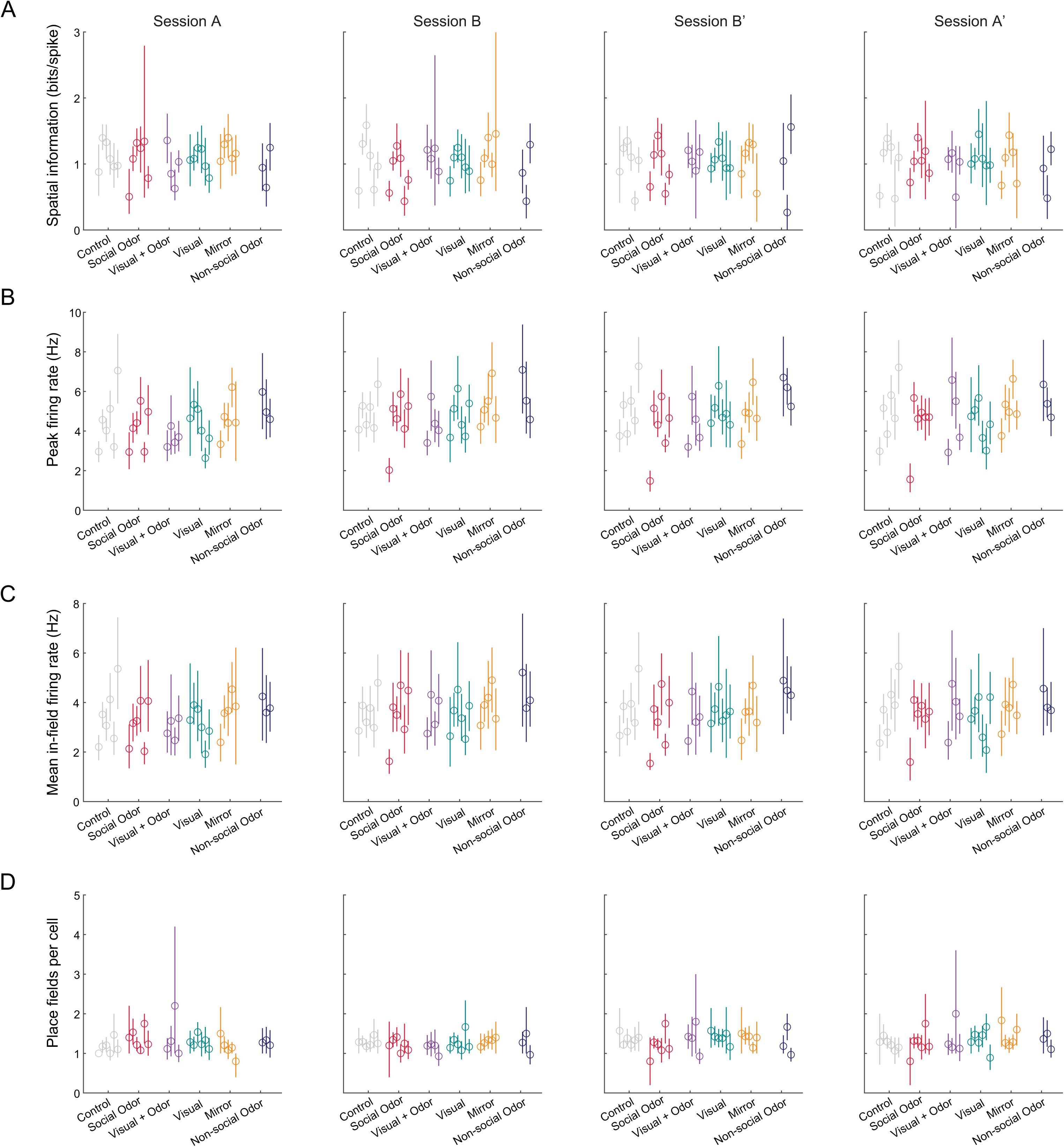
CA2 place cell properties across sessions and conditions. (A) CA2 place cells conveyed similar spatial information across conditions. (B-C) CA2 place cell peak firing rates (B) and in-field firing rates (C) were similar across conditions. (D) The number of place fields per CA2 cell was similar across conditions.

## Notes

### Competing Interest Statement

The authors have declared no competing interest.

### Summary of Updates

Figures 1, 3, 4, 5 updated to include data from a non-social odor condition; Supplemental figures added; Materials and Methods section updated to explain new analyses in updated manuscript. Minor updates to Results and Discussion sections.

